# Parkinson’s disease: a systemic inflammatory disease accompanied by bacterial inflammagens

**DOI:** 10.1101/646307

**Authors:** Büin Adams, J Massimo Nunes, Martin J Page, Timothy Roberts, Jonathan Carr, Theo A Nell, Douglas B Kell, Etheresia Pretorius

**Author notes:** **Corresponding authors: Etheresia Pretorius**, Department of Physiological Sciences, Stellenbosch University, Private Bag X1 MATIELAND, 7602, SOUTH AFRICA, http://www.resiapretorius.net/, **Douglas B. Kell**, Deptartment of Biochemistry, Institute of Integrative Biology, Faculty of Health and Life Sciences, University of Liverpool, Crown St, Liverpool L69 7ZB, UK.

## Abstract

Parkinson’s disease (PD) is a well-known neurodegenerative disease. Recently, the role of gingipains from *Porphyromonas gingivalis* was implicated in Alzheimer’s disease. Here we present evidence of systemic inflammation, accompanied by hypercoagulation; we also show that ginipains from *P. gingivalis* and its LPS may foster abnormal clotting, and that ginipains are present in PD blood, and thus that ginipain’s action on blood may be relevant to PD pathology. Bloods from both PD and healthy blood samples were analysed using thromboelastography (TEG), confocal and electron microscopies, and for cytokine and other circulating biomarkers. We also probed PD and healthy plasma clots with a polyclonal antibody for the bacterial protease, gingipain R1, from *P. gingivalis*. Low concentrations of recombinant gingipain R1 were also added to purified fluorescent fibrinogen. TEG, fibrin(ogen) amyloid formation and platelet ultrastructure analysis confirmed profound hypercoagulation, while the biomarker analysis confirmed significantly increased levels of circulating proinflammatory cytokines. We provide evidence for the presence of the protease, gingipain R1 in PD blood, implicating inflammatory microbial cell wall products in PD.

## INTRODUCTION

Parkinson’s disease (PD) is a neurodegenerative disease caused by the death of dopaminergic neurons in the substantia nigra pars compacta (SNpc), resulting in dopamine deficiency within the basal ganglia. This can lead to a movement disorder with classical parkinsonian motor symptoms, as well as other symptoms. Although a number of Park genes have been identified (Funke et al., 2013), 90% of Parkinson’s disease cases have no identifiable genetic cause (Klein and Westenberger, 2012; Ascherio and Schwarzschild, 2016). PD has a multitude of pathologies (Fujita et al., 2014), ranging from mis-folding of alpha-synuclein to neuro-inflammation, mitochondrial dysfunction, and neurotransmitter-driven alteration of brain neuronal networks (Titova et al., 2017); it also affects all levels of the brain-gut axis (Mulak and Bonaz, 2015).

Neuro-inflammation is an important and well-known feature of PD pathology (More et al., 2013; Nolan et al., 2013; Taylor et al., 2013), and converging evidence further supports the roles of (systemic) inflammation, oxidative stress (Kalia and Lang, 2015) and gut dysbiosis, although the mechanistic details and their full roles in PD pathogenesis are yet to be comprehensively elucidated. It is also noted that there are higher levels of proinflammatory cytokines in brains of PD patients, and inflammation is thought to be a major contributor to the neurodegeneration (Reale et al., 2009). See Figure 1 for an explanatory overview of PD aetiology and our interpretation of the role of systemic inflammation and (hyper)coagulation in this condition. PD patients suffer from a plethora of other (inflammatory) comorbidities (Kell and Pretorius, 2018a), and both vascular risk (Cheng et al., 2017) and cardiovascular autonomic dysfunction are associated with arterial stiffness in these individuals (Kim et al., 2017). Furthermore, heart disease is also associated with dementia in PD (Pilotto et al., 2016). While the interplay between inflammation and neuronal dysfunction is complex, there is mounting evidence that chronic inflammation (Pretorius et al., 2014) with the accompanying dysregulation of circulating inflammatory molecules and the innate immune response, play prominent roles in PD (Kannarkat et al., 2013). It is also becoming recognised that peripheral, as well as brain inflammation, contribute to the onset and progression of the neurodegenerative processes seen in PD (Deleidi and Gasser, 2013; More et al., 2013; Nolan et al., 2013; Taylor et al., 2013; Filiou et al., 2014; Pessoa Rocha et al., 2014).

**Figure 1:**
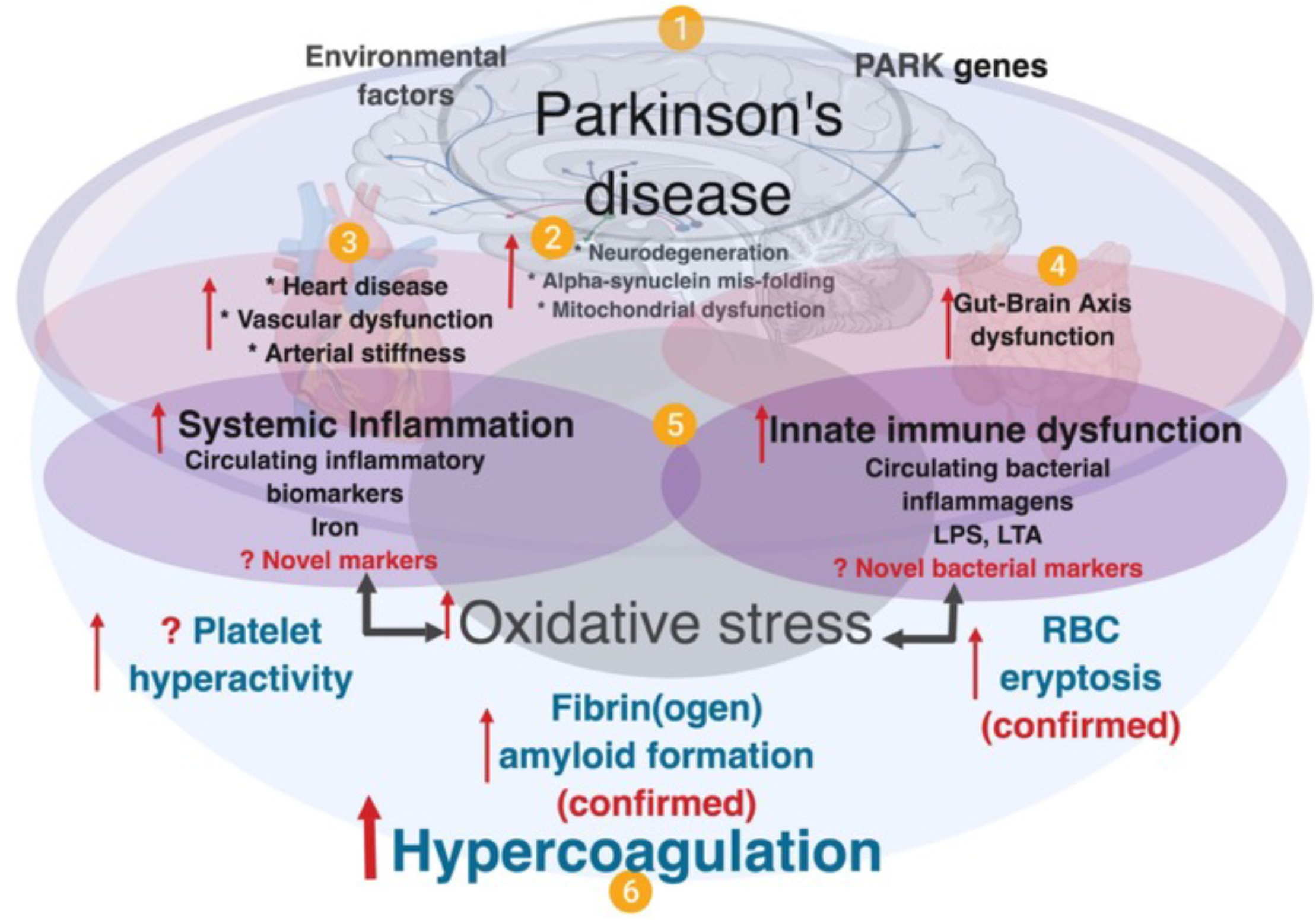
A simplified diagram showing contributing factors in systemic inflammation and hypercoagulation in Parkinson’s disease. **(1)** Parkinson’s Disease is characterized by the presence of PARK genes, and driven by environmental factors with **(2)**, neurodegeneration, and accompanied by **(3)** heart and vascular dysfunction, and also **(4)** gut-brain dysfunction. In PD, dysregulated inflammatory biomarkers and increased circulating bacterial inflammagens (e.g. LPS and LTA), point to **(5)** the presence of systemic inflammation and a dysfunctional innate immune system. Systemic inflammation is usually accompanied by oxidative stress that typically causes a general hypercoagulable state **(6)**, visible as platelet hyperactivity, RBC eryptosis and fibrin(ogen) amyloid formation. Diagram created using BioRender (https://biorender.com/).

Evidence of systemic inflammation in PD includes the presence of increased levels of circulating cytokines such as IL-1β IL-2, IL-10, IL-6, IL-4, TNF-α, C-reactive protein, RANTES and interferon-gamma (INF-γ) (Brodacki et al., 2008; Qin et al., 2016). These markers are accompanied by oxidative stress and might even provide early diagnosis of PD (Lotankar et al., 2017). In addition to dysregulated circulating inflammatory molecules, one of the known hallmarks of systemic inflammation is hypercoagulability, or abnormal clotting potential. In PD, changes in the normal clotting of blood have been described (Sato et al., 2003; Rosenbaum et al., 2013; Pretorius et al., 2014; Infante et al., 2016; de Waal et al., 2018; Pretorius et al., 2018c). Most of these circulating inflammatory biomarkers act as ligands to receptors on platelets (Olumuyiwa-Akeredolu et al., 2019), resulting in downstream signaling events with accompanying platelet hyperactivity and aggregation. RBCs also become eryptotic (programmed cell death in RBCs) due to ligand binding and oxidative stress (Pretorius et al., 2014).

What is not immediately clear is the actual origin of the inflammation and how and why it is chronic. For this and other diseases (Potgieter et al., 2015; Kell and Kenny, 2016; Pretorius et al., 2016a; Pretorius et al., 2017a; de Waal et al., 2018; Kell and Pretorius, 2018a; Kell and Pretorius, 2018b) we have brought together evidence that a chief cause may be (dormant) microbes that upon stimulation, especially with unliganded iron (Kell, 2009), can briefly replicate and shed potent (and well known) inflammagens such as lipopolysaccharide (LPS) and lipoteichoic acid (LTA) (Kell and Pretorius, 2015; Kell and Pretorius, 2018a). These are well-known ligands for receptors such as Toll-like receptor 4 (TLR4) and can thus stimulate inflammation, as observed through a variety of inflammatory cytokines (Olumuyiwa-Akeredolu et al., 2019).

Another set of (novel) bacterial inflammagens that might cause damage to fibrin(ogen) proteins when present in circulation is represented by a group of proteases synthesized by *Porphyromonas gingivalis*. *P. gingivalis* is a Gram-negative anaerobic bacterium that is deemed a keystone pathogen in the oral cavity with the capacity to shift symbiotic homeostasis into a dysbiotic state characteristic of periodontal pathogenesis (Darveau et al., 2012; How et al., 2016). Accordingly, this bacterium is significantly associated with and demonstrated to be a cause and driver of chronic periodontitis – the most common oral disorder among adults (Nazir, 2017). The bacterium’s entry into the circulation has been well-documented (Silver et al., 1977; Parahitiyawa et al., 2009; Tomas et al., 2012; Ambrosio et al., 2019); and it enters through teeth care activities and oral ulcerations. Periodontal pathologies are known to be linked to systemic inflammation (Hajishengallis, 2015; Leira et al., 2018; Torrungruang et al., 2018), and *P. gingivalis* in particular, is associated with a cohort of diseases including non-insulin dependent diabetes mellitus (Makiura et al., 2008; Blasco-Baque et al., 2017), Alzheimer’s disease (Singhrao et al., 2015; Dominy et al., 2019), rheumatoid arthritis (Okada et al., 2013; Mikuls et al., 2014; Jung et al., 2017), cardiovascular disease (Deshpande et al., 1998; Aarabi et al., 2015; Chistiakov et al., 2016; Leira et al., 2018) and atherosclerotic vascular tissue (Deshpande et al., 1998; Velsko et al., 2014; Olsen and Progulske-Fox, 2015).

The bacterium uses oligopeptides as its main nutrients, that it obtains via protease activities. Recently, emphasis was placed on both the bacterium and its group of endogenous cysteine proteases called gingipains, in developing Alzheimer’s disease, where gingipains were implicated in disease causation and suggested as possible disease intervention targets (Dominy et al., 2019). Gingipains are important protease of *P. gingivalis* and their proteolytic activity plays an important part of the functioning of the bacterium, as it is essential for obtaining nutrients via protein degradation, adherence to host surfaces and further colonisation (Guo et al., 2010). Gingipains are also known to play an important role in neutralizing the host defences by degrading of antibacterial peptides (Guo et al., 2010), and interfering or evading the host complement system (Slaney and Curtis, 2008). These enzymes cleave proteins at the C-terminal after arginine or lysine residues and are classified accordingly: gingipain R is arginine-specific and gingipain K is lysine-specific.

There are two types of arginine-specific gingipains: RgpA which seems to be the more virulent (Imamura et al., 2000) and RgpB. Not only are gingipains found on the cell surface of *P. gingivalis*, but are also secreted from the bacterium and can thus enter the circulation, where it may interact with various circulating blood proteins, including clotting proteins. Studies have demonstrated fibrinogen-adhesive and fibrinogenolytic effects arising from each gingipain type (Lantz et al., 1986; Imamura et al., 1995; Pike et al., 1996; Ally et al., 2003). Further, the effect of gingipain proteases on fibrinogen increases the propensity for bleeding at periodontal sites (symptom of periodontitis) thereby enabling *P. gingivalis* access to nutrient sources (heme-containing proteins) and inadvertently the circulation. The interference of these proteases in coagulation may not be exclusive to fibrinogen, and interactions have been shown with factor IX prothrombin (Imamura et al., 2001), factor X (Imamura et al., 1997) and prothrombin (Imamura et al., 2001), as well as the stimulation of the kallikrein/kinin pathway (Imamura et al., 1994). Since periodontitis disposes an individual to an exaggerated risk of developing Parkinson’s disease (Kaur et al., 2016; Chen et al., 2017; Chen et al., 2018) and because the activity of *P. gingivalis* and gingipains have recently been highlighted in Alzheimer’s patients (Dominy et al., 2019), we might expect the presence of this bacterium and its molecular products (e.g. proteases and LPS) to be found in the circulation of PD individuals too.

In this paper, we therefore aim to offer further evidence of the significant role of systemic inflammation and circulating inflammagens in the development of PD. Here we show the extent of the dysregulated systemic inflammatory biomarker profile, hypercoagulability and particularly platelet hyperactivity in PD patients compared to healthy individuals, and how dysregulated inflammatory circulating molecules could, in part, be responsible for blood hypercoagulability and platelet dysfunction. We also study whole blood clot formation using thromboelastography, and look at platelet ultrastructure using scanning electron microscopy. Furthermore, we hypothesize how these dysregulated inflammatory molecules might act as ligands when they bind to platelet receptors, resulting in activation of platelet signaling cascades. We argue that the levels of inflammatory molecule dysregulation point to innate immune system activation, which is supportive of our previous published results regarding the presence of LPS in/near hypercoagulated blood clots (de Waal et al., 2018). We confirm the presence of amyloid fibrin(ogen) in the current sample, using amyloid-specific markers (previously we used only thioflavin T as a marker of aberrant clotting in PD (Pretorius et al., 2018a)). To date, *P. gingivalis* and its molecular signatures are yet to be discovered in PD tissue (other than the oral cavity). We present evidence (using fluorescent antibodies against gingipains), that members of the gingipain protease family are present in clots from PD samples, but not significantly present in healthy plasma clots. We also add purified RgpA to purified fibrinogen marked with a fluorescent Alexa 488 marker, and show how it potentially can hydrolyze fibrinogen proteins and that gingivalis LPS may act together with gingipains to foster aberrant clot formation (see supplementary Figure A for a layout of our experiments).

## MATERIALS AND METHODS

### Ethical clearance and consent

Ethical clearance was received for this study from the Health Research Ethics Committee (HREC) of Stellenbosch University (South Africa) (approval number HREC Reference #: S18/03/054) and the Health Department of Western Cape research number (WC_201805_023). Written informed consent was obtained from all participants followed by whole blood sampling in citrated tubes. All participants received a unique number that was used to guarantee discretion throughout this study. All investigators were certified in Good Clinical Practice and ethical codes of conduct.

### Study design, setting and study population

A cross-sectional design was followed in collaboration with a neurologist, who provided whole blood (WB) from Parkinson’s Disease (PD) patients at Tygerberg Hospital in the Western Cape. Whole blood from healthy controls was collected by a Health Professions Council of South Africa (HPCSA) registered Medical Biological Scientist and phlebotomist (MW: 0010782) at the Department of Physiological Sciences, Stellenbosch University. A total of n=81 volunteers were included (n=41 healthy controls, and n=40 PD patients) as part of the study population. PD volunteers were recruited with the following inclusion criteria: (i) a confirmed diagnosis by a neurologist and the diagnosis of these patients will included the use of the Unified Parkinson’s Disease Rating Scale (UPDRS), as well as the Hoehn and Yahr scale used to rate the relative level of the PD disability, (ii) males and females of any age, (iii) not on any anticoagulant medication. Participants who were unable to provide written consent were omitted from this study. To limit and exclude confounding factors, both healthy and PD volunteers were only included if they were not diagnosed with tuberculosis, HIV or any malignancies. The inclusion criteria for healthy age-matched volunteers included were also: (i) no use of chronic medication (ii) no prior history of thrombotic disease or inflammatory conditions (iii) non-smokers, (iv) not on any chronic antiplatelet therapy/ anticoagulant medication or any contraceptive/hormone replacement therapy (v) were not pregnant and/or lactating. PD is a progressive condition which tend to evolve from mild unilateral symptoms through to end-stage non-ambulatory state. See supplementary Table 1 and 2 for the milestones in the illness as accurately outlined in the Hoehn and Yahr staging system.

### Collection of whole blood (WB) and preparation of platelet poor plasma (PPP) samples from healthy controls and PD patients

Whole blood from PD patients and heathy controls were collected by sterile sampling techniques in citrate and ethylenediaminetetraacetic acid (EDTA) tubes, as well as serum separating tubes (SST) that were kept at room temperature (∼22°C) for 30 min. Platelet poor plasma (PPP) was prepared from citrate tubes that were centrifuged at 3000 x *g* for ten minutes at room temperature (∼22°C). The PPP was then aliquoted into labelled 1.5 mL Eppendorf tubes, and stored at −80°C until cytokine analysis. EDTA whole blood and SST were analysed by the local PathCare laboratory (Stellenbosch) for glycosylated haemoglobin (HbA1c), total cholesterol (TC), low-density lipoprotein cholesterol (LDL-c), high-density lipoprotein cholesterol (HDL-c), triglyceride (TG) and non-high-density lipoprotein (non-HDL); TC/HDL ratio was calculated as a marker of cardiovascular risk.

### Thromboelastography (TEG) of whole blood (WB)

Clot kinetics/property analysis was completed by means of a Thromboelastography (TEG) (Thromboelastograph 5000 Hemostasis Analyzer System, Haemonetics S.A. Signy-Avenex, Switzerland), on both control and PD WB samples. 340 μL of WB samples were placed in a disposable TEG cup to which 20 μL of 0.2 mol/L CaCl_2_ was added. CaCl_2_ is necessary to reverse the effect of the sodium citrate anticoagulant in the collection tube (i.e. recalcification of blood) and consequently initiate coagulation.

### Scanning electron microscopy of whole blood (WB) smears

WB smears were prepared by placing 10µL WB of each of the samples on cover slips. Samples were washed with Gibco™ PBS (phosphate-buffered saline), pH 7.4 (ThermoFisher Scientific, 11594516) before fixing with 4% paraformaldehyde for a minimum of 30 minutes. Once fixed, samples were washed 3 × 3 minute with PBS followed by a second 30-minute fixation step in 1% osmium tetroxide (Sigma-Aldrich, 75632). A final 3 × 3 minute PBS wash step was performed before samples were serially dehydrated in ethanol with a final 30-minute dehydration step using hexamethyldisilazane (HMDS) ReagentPlus® (Sigma-Aldrich, 379212). Samples were then carbon coated before being imaged on Zeiss MERLIN™ field emission scanning microscope and micrographs were captured using the high resolution InLens capabilities at 1 kV.

### 20-Plex cytokine analysis using platelet poor plasma (PPP)

Stored PPP samples of PD (n= 40) and healthy controls (n=41) participants were transferred from −80°C to −20°C 24 hours preceding the multiplex analysis. The samples were then analysed in duplicate by means of Invitrogen’s Inflammation 20-Plex Human ProcartaPlex™ Panel (#EPX200-12185-901) and read on the Bio-Plex^®^ 200 system (Bio-Rad, 2016). The data is expressed in pg⋅mL^−1^. 20 anti- and pro-inflammatory molecules were measured in a multiplex analysis and biomarkers measured included 4 anti-inflammatory molecules (IFN-α, IL-4, IL-10, IL-13), and 16 pro-inflammatory molecules (for the full list of cytokines, see the table in results section).

### Recombinant gingipain R1 protease (RgpA) and Gingipain R1 antibody

Platelet poor plasma (PPP) was used to prepare clots from PPP from 6 healthy and 10 PD samples. Thrombin was donated by the South African National Blood Service; it was solubilized in PBS containing 0.2% human serum albumin to obtain a concentration of 20 U⋅mL^−1^ and was used at a 1:2 ratio to create extensive fibrin networks. This was followed by fixation with 10% neutral buffered formalin (NBF). After phosphate-buffered saline (PBS) (pH=7.4) washing steps, samples were blocked with 5% goat serum (in PBS), and incubated with gingipain R1 polyclonal antibody (Abbexa, abx 107767) (1:100 in 5% goat serum) for one hour at room temperature in the dark. The samples were finally washed and a coverslip was mounted with a drop of Dako fluorescence mounting medium on a microscopy slide for confocal analysis. The prepared samples were viewed on a Zeiss LSM 780 with ELYRA PS1 confocal microscope using a Plan-Apochromat 63x/1.4 Oil DIC objective. The gingipain R1 FITC antibody was excited at 488 nm, with emission measured at 508 to 570 nm. As a positive control, we also incubated an exogenous aliquot of the protease, recombinant gingipain R1 protease (RgpA), with healthy PPP for 30 minutes, followed by exposure to its fluorescent antibody. RgpA (Abcam. ab225982) was added at a final concentration of 500 ng.L^−1^.

### Recombinant Gingipain R1 protease and Alexa 488-conjuagted purified fibrinogen

Purified (human) fibrinogen with Alexa 488 (ThermoFisher: F13191) was prepared to a final concentration of 2 mg⋅mL^−1^. Clots (with and without the protease, RgpA) were prepared by adding human thrombin as per the above protocol. Clots were also viewed with the confocal microscope and fluorescent fibrinogen was excited at 488 nm, with emission measured at 508 to 570 nm. As the gingipains antibody used above has the same excitation and emission as the purified fibrinogen, we could not trace the added gingipains with this antibody. We also incubated purified fluorescent fibrinogen with LPS from *P. gingivalis* (10ng.L^−1^) with and without RgpA (both 100 and 500 ng.L^−1^). Where we combined the LPS and the RgpA, we added it simultaneously and the incubation period was also 30 minutes.

### Confocal analysis of plasma clots to show amyloid fibrin(ogen)

Three fluorescent amyloid markers were added to control and PD PPP to illuminate amyloid protein structure, and were used as follows: 5 µM thioflavin T (ThT), 0.1 µL (stock concentration as supplied) of AmyTracker 480 and 0.1 µL (stock concentration as supplied) of AmyTracker 680 were added to the sample to incubate for 30 minutes. A working solution of AmyTracker was made in PBS at a 1:20 ratio. Control and PD PPP clots were prepared by adding thrombin to activate fibrinogen and create extensive fibrin fibre networks. Using the same microscope and objective as above, three channels were setup to visualise the amyloid markers. Amytracker 480 was excited by the 405 nm laser, with emission measured at 478 to 539 nm; Amytracker 680 was excited by the 561 nm laser, with emission measured at 597 to 695 nm; and ThT was excited by the 488 nm laser, with emission measured at 508 to 570 nm. ThT may also be excited by the 405 laser, and has a wide spectra where fluorescence can be detected (Sulatskaya et al., 2017). We allowed these two stains, which both target amyloid structures, to overlap in the microscope setup to produce a combination blue channel of amyloid signal, alongside the isolated signal from Amytracker 680 in the red channel and ThT in the green channel (Page et al., 2019). Micrographs of the prepared clots were captured as 3×3 and 2×2 tile images, and 75 images from 25 PD patients and 39 images from 9 control donors were acquired. Gain settings were kept constant for all data acquisition and used for statistical analyses; however, brightness and contrast were adjusted for figure preparation. The mean and the standard deviation from the histogram of each image were recorded and used to calculate the coefficient of variance (CV), which is defined as SD ÷ mean. This metric was used to quantify and discriminate the signal present between control and PD PPP clots. CVs of the control and disease group were compared by the Mann-Whitney test in GraphPad Prism 7.04 with significance accepted at p<0.05.

### Statistical analyses

Statistical analysis was performed using R version 3.5.1 with specific packages detailed below. Three variations of logistic regression modelling were investigated to determine the strength of association between measured parameters and Parkinson’s disease status. For all three models, Wald p-values are reported in a manner that allows inter-model comparison. Logistic regression, based on the glm from the built-in stats package, was performed between Parkinson’s status (binary) and all individual parameters both with no adjustment (Model 1) and with adjustment for age and gender (Model 2). Ordinal logistic regression, based on the clm method in the ordinal package, was performed between the Hoehn and Yahr severity scale and all individual parameters without adjustment due to sample size requirements (Model 3). Mann Whitney non-parametric tests were also performed and contrasted with the results from logistic regression. Although the Mann Whitney test was found to be more sensitive (identifying a super-set of parameters as significant), upon inspection of the populations trends and outliers, the logistic regression model was deemed more appropriate due to (a) better aligned with the goal of identification of regressive trends, (b) being more *conservative*, especially in the presence of significant outliers and (c) easily extended to adjustment and ordinal modelling scenarios. PCA analysis was performed using the prcomp method from the built-in stats package. Our raw data files are accessible at: https://1drv.ms/f/s!AgoCOmY3bkKHibs-vg0EUq3N5SogfA.

## RESULTS

Tables 1 shows summary statistics of markers from WB for healthy and PD populations along with statistical significance values between the populations for all three regression models. More specifically, the first part of Table 1 shows the 7 TEG clotting parameters as well as lipid profile, HbA1c and ultrasensitive CRP levels. The second part of Table 1 shows anti-inflammatory and pro-inflammatory cytokine markers.

**Table 1:**
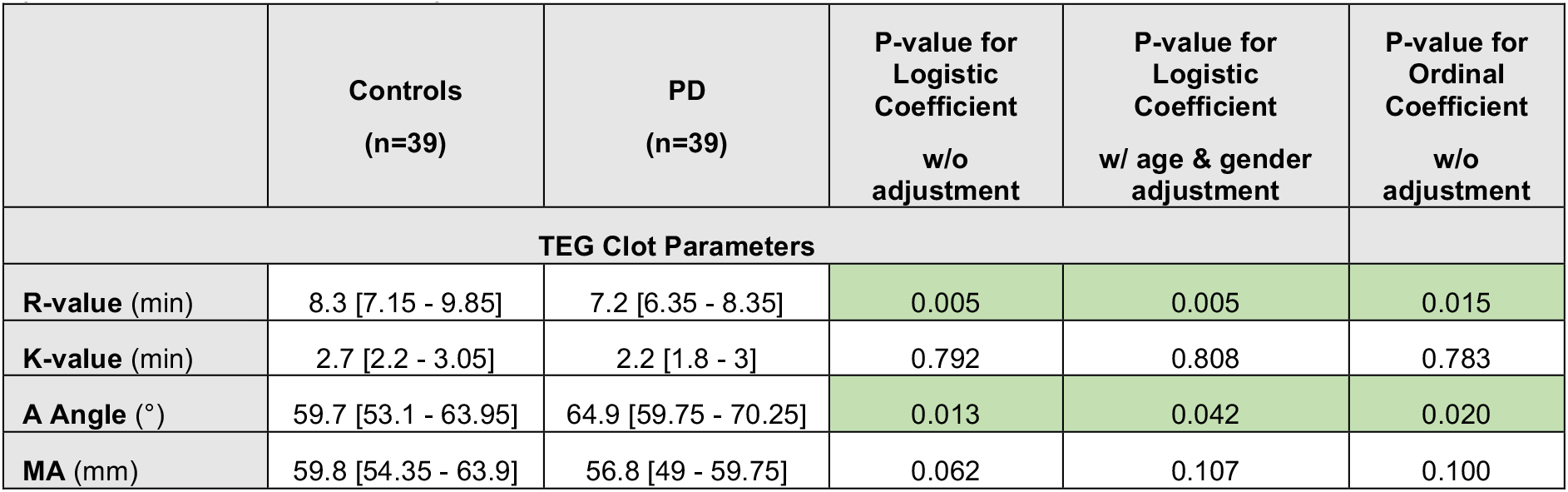
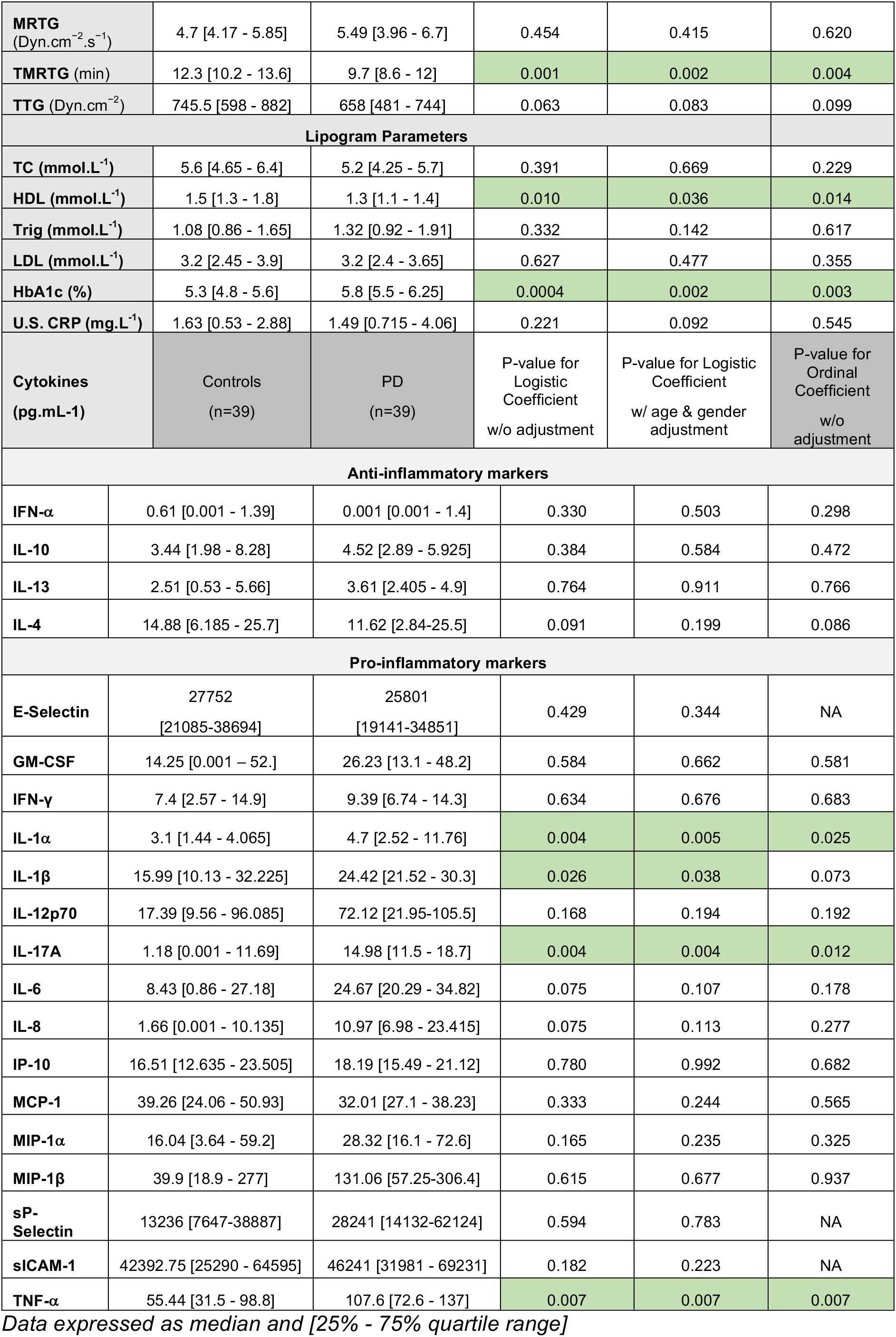
Thromboelastography results showing seven viscoelastic parameters assessing coagulation properties of healthy control and Parkinson’s disease WB samples. Whole blood lipid profiles, HbA1c and ultrasensitive CRP as well as anti-inflammatory and pro-inflammatory cytokine profiles of healthy and PD volunteers are also shown.

The three regression models consistently identify the same parameters as significant (at level of 0.05) in most cases. The exception is IL-1β which was not significantly predictive in the ordinal logistic regression model (i.e. not predictive of the scale of the disease). One can also observe that significant differences exist across all groupings except anti-inflammatory markers. To summarize, the following parameters in each group can be identified as significantly different:

- **TEG parameters**: R, Angle, TMRTG
- **Lipogram parameters**: HbA1c, HDL
- **Anti-inflammatory markers**: None
- **Pro-inflammatory markers**: IL-1α, IL-17A, TNF-α, IL-1β

Figure 2 shows box and whisker plots of these parameters, illustrating population differences and the presence of significant outliers. Figure 3 shows a lattice of parameter cross-plots along with correlation coefficients in the upper diagonal. One can observe correlations between the R, TMRTG and A Angle parameters. (Supplementary Figure B shows a visualization based on PCA analysis of the combined data). Ellipses for Parkinson’s disease status are overlaid but were not part of the analysis. Notice that the first two principal components capture around 30% of the variance in the data.

**Figure 2:**
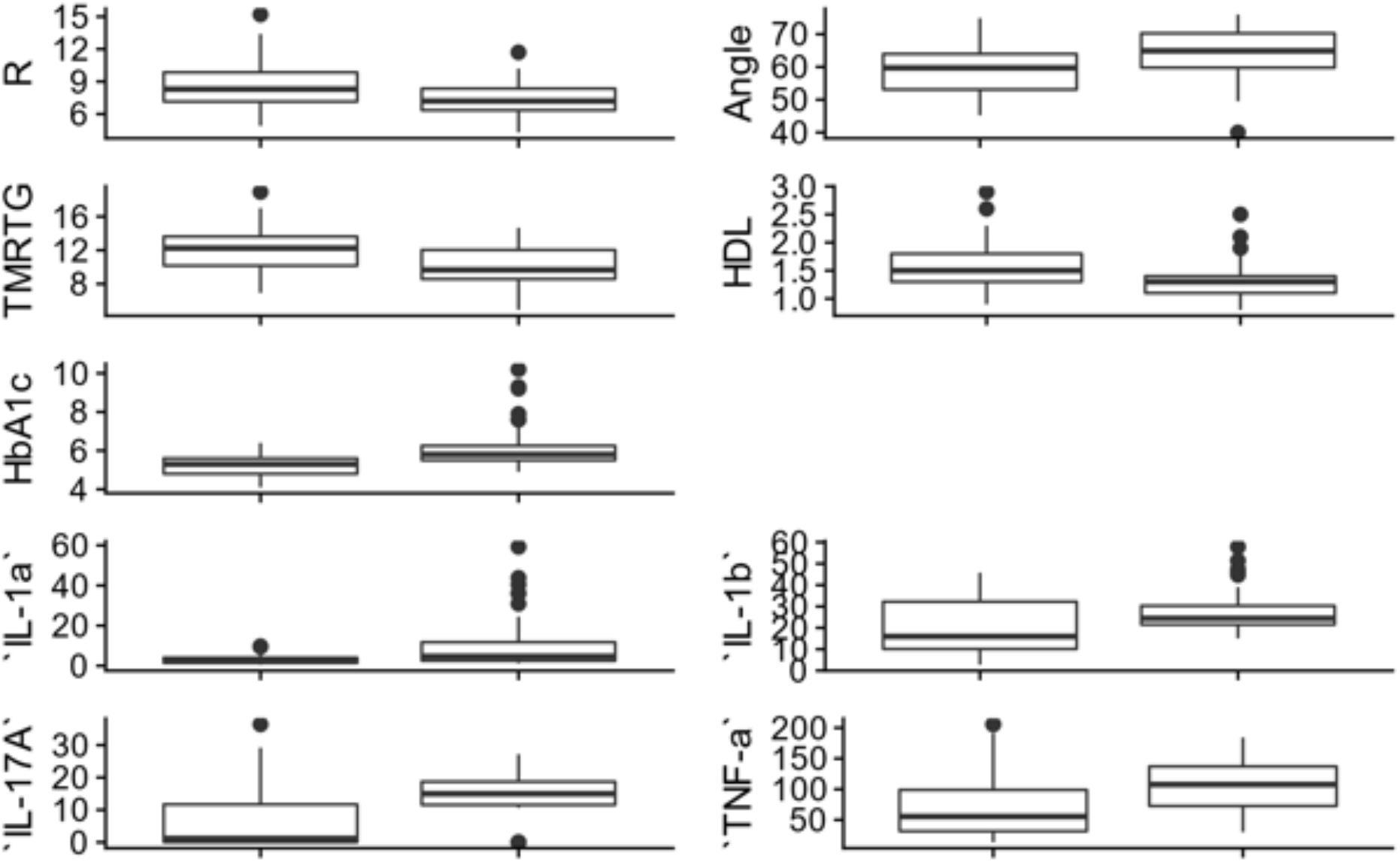
Box and whisker plots showing the distribution of parameters for control (left box) and PD (right box) populations for parameters determined to be significantly different.

**Figure 3:**
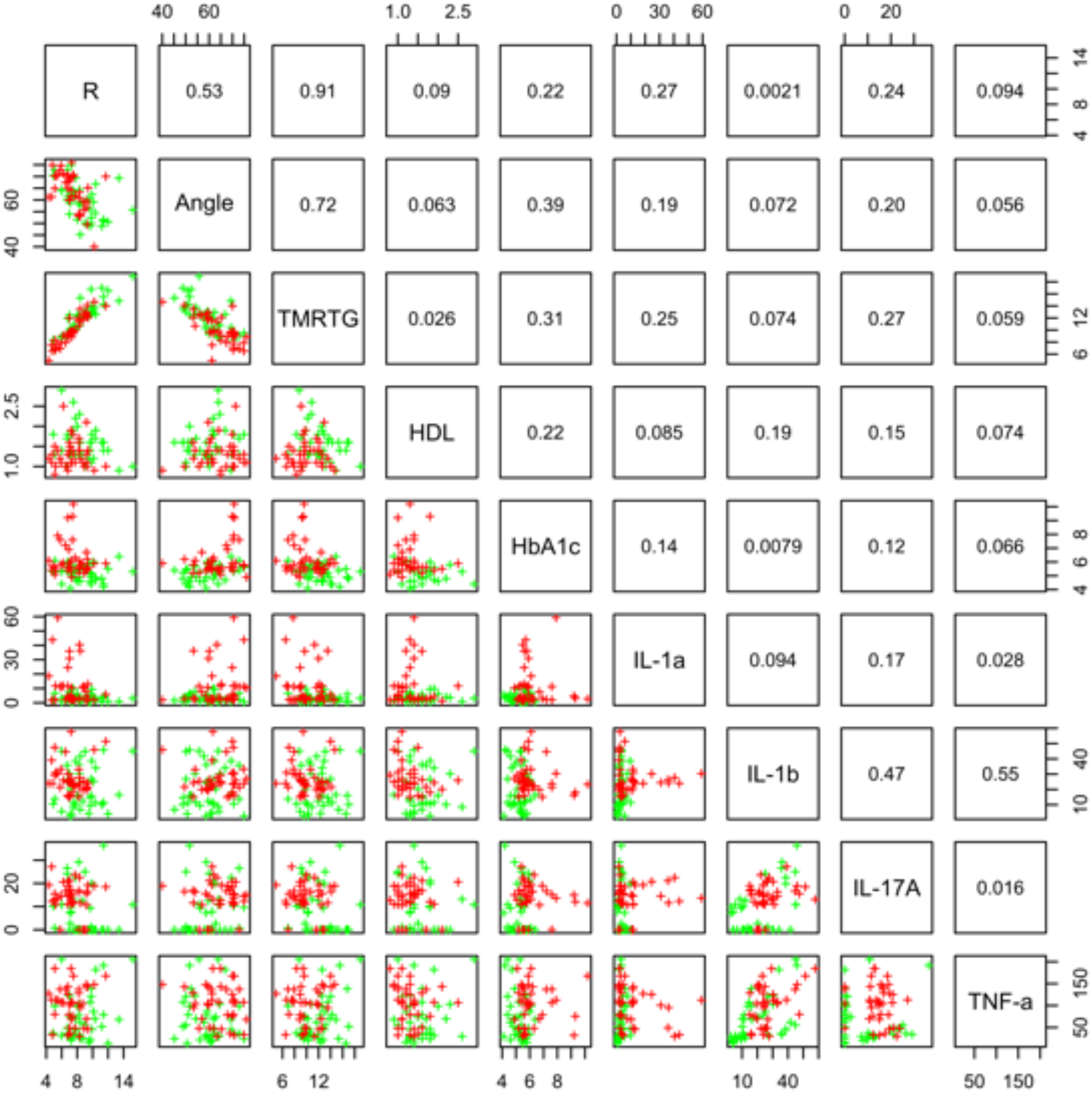
Lattice of cross-plots of statistically significant parameters colored by Parkinson’s disease status (Green = Control). The upper diagonal shows correlation coefficients.

### Thromboelastography, cholesterol and HbA1c levels, and ultrasensitive CRP

HbA1c levels were significantly increased in our PD sample and a slightly dysregulated lipid profile was also noted (see Table 1). TEG results point to the fact that PD WB is hypercoagulable. TEG analysis exhibited significant differences in five of the groups of the assessed parameters. The PD group presented a significant increase in the initial rate of clot formation (R-value). Significant elevation in alpha angle (A angle) suggests more cross-linking of fibrin fibres, and time to maximum rate of thrombus generation (TMRTG) was decreased. These results have significance to our RgpA results that we discuss later.

### Scanning electron microscopy of whole blood

Figure 4 shows representative SEM micrographs of platelets seen in WB smears. SEM analysis of WB smears, of healthy individuals usually show platelets as roundish cellular structures, with only slight pseudopodia formation due to contact activation with glass cover slips. This has also previously been noted in various publications (Page et al., 2018; Pretorius et al., 2018d; Page et al., 2019). In the PD sample, platelets showed substantial (hyper)activation, spreading, as well as aggregation (Figure 4).

**Figure 4:**
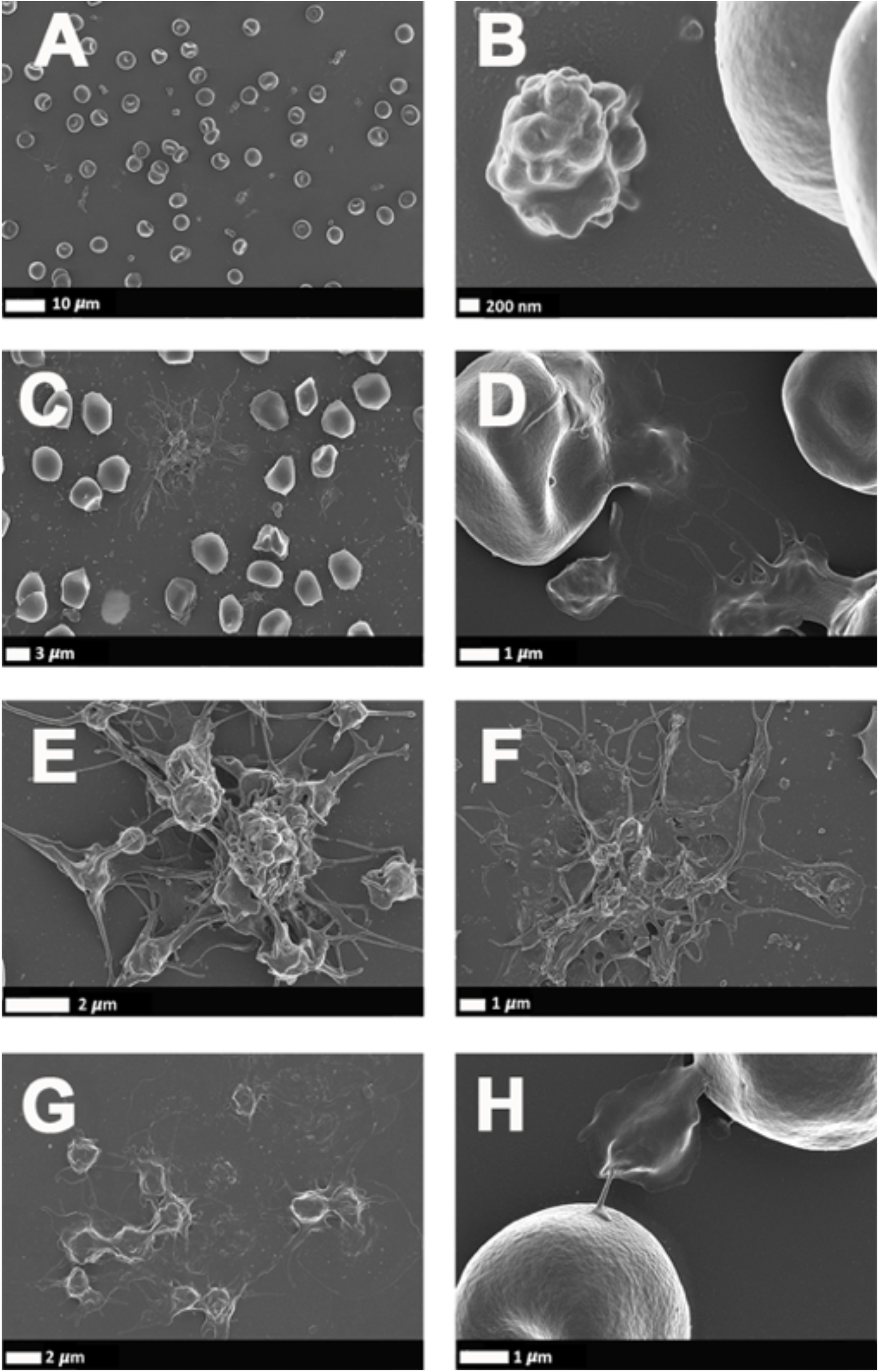
**(A and B)** Scanning electron microscopy of whole blood smears showing representative platelets from healthy individuals. (**C to H)** Whole blood smears from Parkinson’s disease (PD) individuals showing hyperactivated platelets. **(C to H)** PD platelets agglutinating to RBCs; **(D and H)** PD platelet spreading **(G)** and PD platelet aggregation **(C, E and F)**.

### The identification of gingipain R1 in Parkinson’s disease blood samples with its fluorescent antibody

For each control and PD sample, we viewed unstained and antibody-stained clots (Figure 5A to F). Unstained clots from both control (Figure 5A) and PD donors (Figure 5C and E) showed no fluorescent signal. Antibody-stained control clots showed little to no detectable fluorescent signal (Figure 5B), whereas PD samples showed substantial fluorescent signal (Figure 5D and F). Thus, our results indicate the presence of RgpA, an arginine-specific variant of virulent gingipain proteases produced by *P. gingivalis* in the blood of PD patients. As positive control, we exposed controls to a tiny concentration of exogenous recombinant RgpA. followed by polyclonal antibody staining against RgpA (Figure 5G). A distinct but minimal signal was now present. This was expected, as the concentration of RgpA added to healthy PPP was very low (500 ng.L^−1^ final exposure).

**Figure 5:**
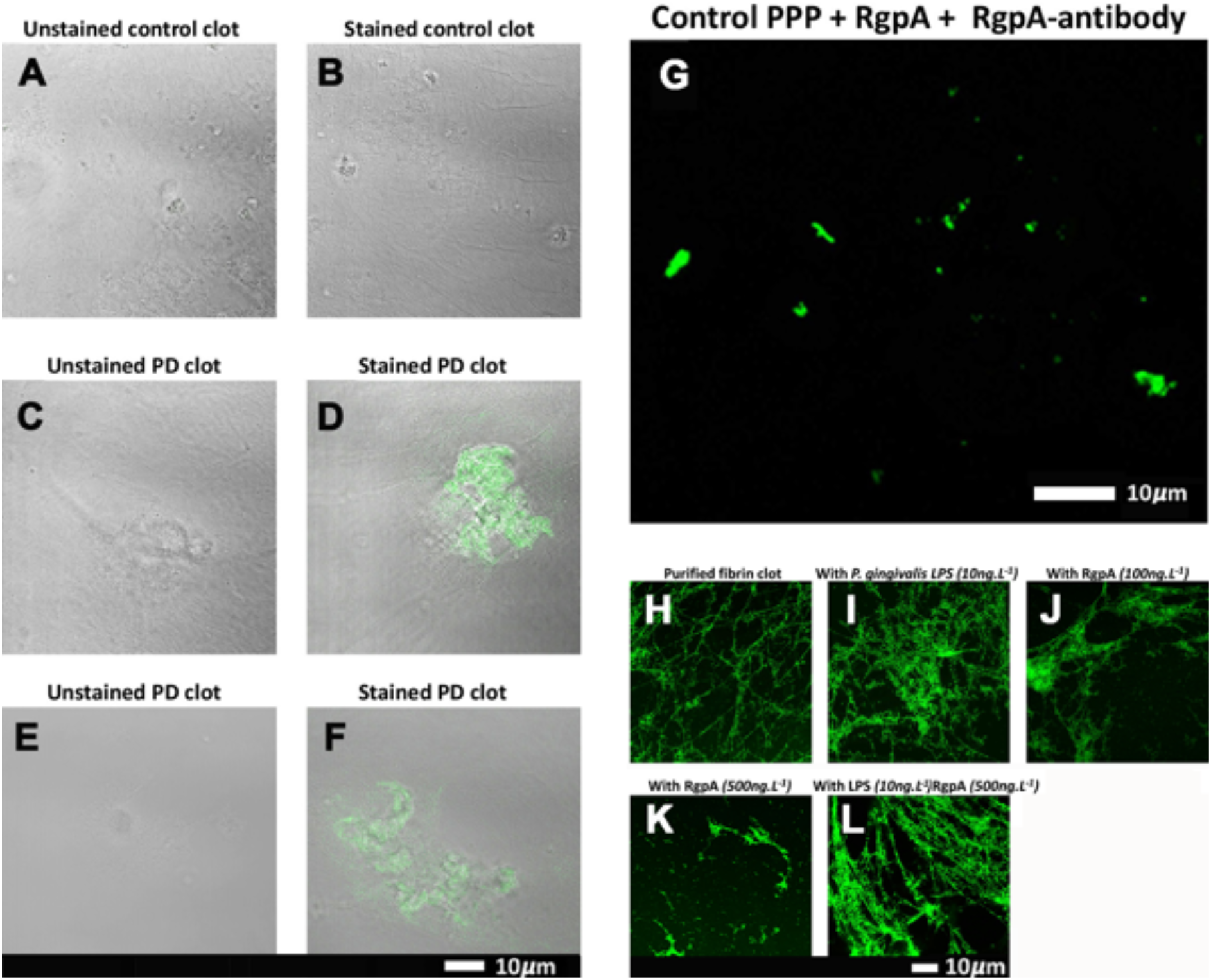
**(A TO G)** Confocal microscopy images of PPP clots stained with the RgpA polyclonal antibody (1:100) from healthy individuals and individuals suffering from Parkinson’s disease. The unstained **(A)** and stained **(B)** control exhibits no fluorescent signal as well as both the unstained Parkinson’s disease PPP clots **(C & E)**. Fluorescent signal of the RgpA antibody is prominently detected in stained Parkinson’s disease PPP clots **(D & F)**. **(G)** represents a positive control in which a control sample that is absent of fluorescent signal received an exogenous load of RgpA. **(H TO L)** Confocal microscopy images of fibrin networks formed from purified fibrinogen (with added Alexa488 fluorophore) incubated with and without RgpA, and LPS from *P. gingivalis*, followed by addition of thrombin to create extensive fibrin(ogen) clots. **(H)** Representative purified fibrin(ogen) clot. **(I)** A representative clot formed after purified fibrinogen was incubated with 10ng.L^−1^ *P. gingivalis* LPS. **(J)** A representative clot formed after purified fibrinogen was incubated with 100ng.L^−1^ RgpA and **(K)** 500ng.L^−1^ RgpA. **(L)** A representative clot after purified fibrinogen was simultaneously exposed to a combination of *P. gingivalis* LPS (10ng.L^−1^) and RgpA (500ng.L^−1^).

### The analysis of clots formed from fibrinogen incubated with recombinant gingipain R1

Confocal microscopy was used to visualize the clot structure of purified fibrin(ogen) marked with Alexa 488, with and without exposure to recombinant gingipain R1 (500 ng.L^−1^), and with and without exposure to *P gingivalis* LPS (Figure 5H to L). Note that fibrinogen was pre-incubated with the inflammagens, followed by clot formation due to the addition of thrombin. Figure 5H is a representative purified fluorescent fibrin(ogen) clot, showing a fibrin network with distinctive fibres. Figure 5I shows a representative fibrin(ogen) clot after fluorescent fibrinogen was incubated with *P. gingivalis* LPS. Fibrin networks display a denser and more matted network. Purified fibrinogen was also exposed to two concentration of RgpA (100ng.L^−1^) (Figure 5J) and 500ng.L^−1^ (Figure 5K). RgpA greatly inhibited fibrin formation synthesis in a concentration-dependent manner. A combination of both the LPS and RgpA (500ng.L^−1^) was also added simultaneously to purified fibrinogen, and the resulting clot is shown in Figure 5L. Interestingly, this clot appeared similar to the clot where only LPS was added (Figure 5I). We suggest that the LPS and the protease might function together, where the protease might hydrolyze the fibrin(ogen) peptides but the LPS might simultaneously cause aberrant coagulation.

### Confocal analysis of plasma clots

Confocal analysis, as well as raw data of the clot analysis are shown in the Supplementary material and in Figure 6. Control and Parkinson’s Disease platelet poor plasma clots, with markers illuminate amyloid fibrin(ogen) protein structure were imaged on a confocal microscope. Control clots display disperse signal. PD samples contain significantly greater amyloid-specific signal than control donors in all three channels: blue (p=0.0002), red (p=0.02) and green (<0.0001).

**Figure 6:**
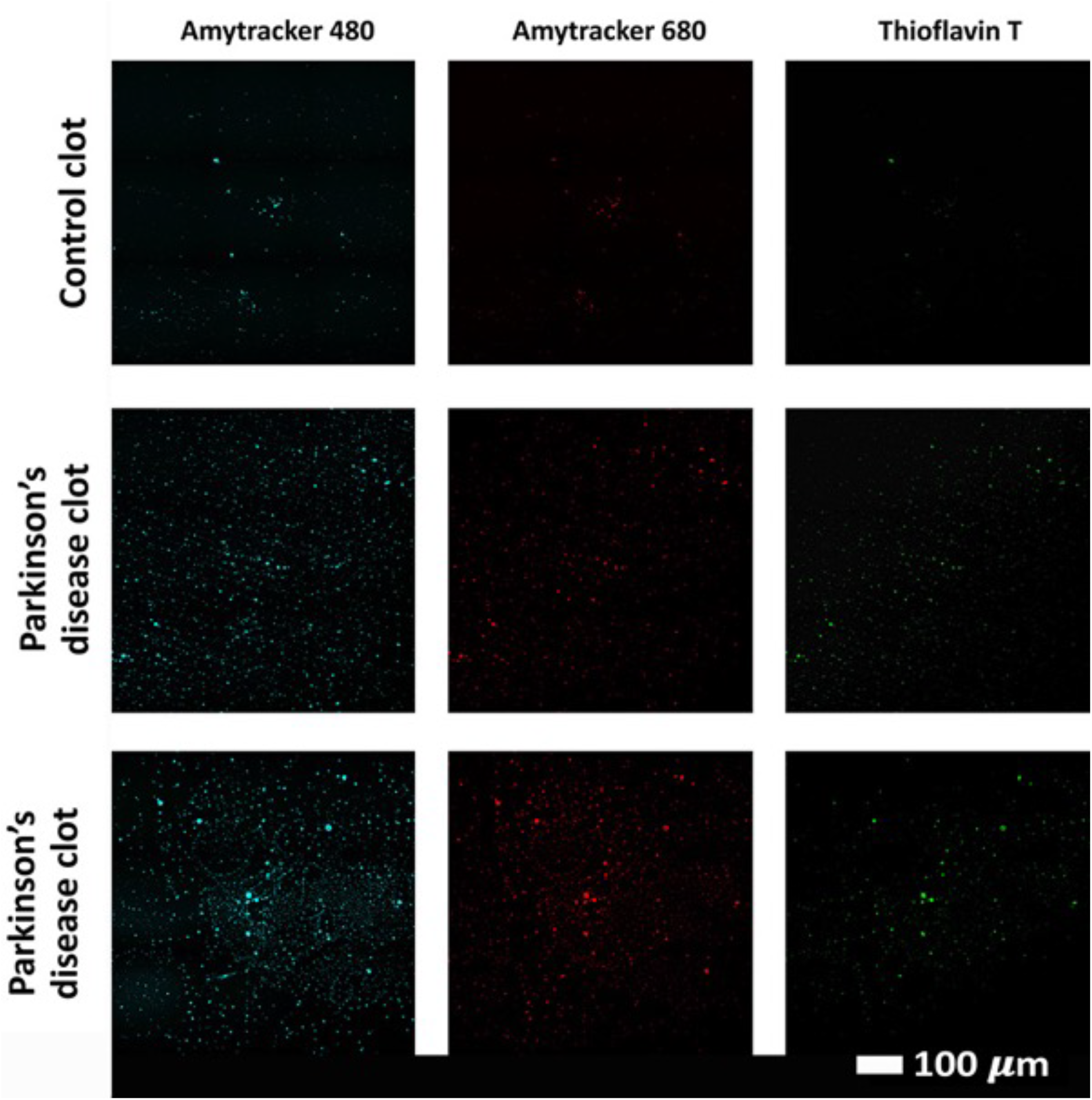
Examples of clots created with platelet poor plasma (PPP) for a representative control and two representative Parkinson’s disease individuals to show amyloid fibrin(ogen) protein structure. Three fluorescent markers that bind amyloid protein were used, Amytracker 480, 680 and Thioflavin T (as previously used for amyloid fibrin structure (Pretorius et al., 2017c; de Waal et al., 2018).

## DISCUSSION

In this paper, we show that in PD, there is a dysregulated systemic inflammatory biomarker profile, and that whole blood of these individuals are hypercoagulable (as seen with our TEG analysis), while platelets are hyperactivated (SEM analysis). The most significant differences were noted in the HbA1c, R-value, Alpha angle, TMRTG (TEG parameters), IL-1α, IL-17A, TNF-α, IL-1β (pro-inflammatory markers) and HDL (note that they were not significantly predictive of PD severity from Hoehn & Yahr). Taken together, these results point to ainter-linked relationship between the hypercoagulability, inflammatory molecule presence, and platelet activation.

The pro-inflammatory profile may relate to blood clotting in various ways. These molecules may all act as outside-in signaling ligands (Durrant et al., 2017) that bind to platelet receptors, followed by inside-out signaling (Faull and Ginsberg, 1996) and ultimately platelet dysfunction. The consequence after inflammatory molecule receptor binding, is platelet activation, visible as platelet (hyper)activation, spreading and aggregation (or clumping). The subsequent platelet pathology, together with other changes in the haematological system such as anomalous fibrin(ogen) protein structure (discussed below) and RBC eryptosis (previously noted (Pretorius et al., 2018c)), all point to inflammation profile of systemic change. Here, the inflammatory molecules in our panel that showed the most significance in PD, and particularly IL-1α, IL-1β, IL-17A, and TNF-are all known to be dysregulated in cardiovascular disease and their presence in circulation might be linked to atherosclerosis (Libby, 2017; Wang et al., 2017).

### Platelet (hyper)activation in Parkinson’s disease and why they might be targets for circulating of cytokines that are increased

We seek to provide a possible explanation for the significant platelet activation that we have noted by closely looking at our cytokine results, and particularly some of the most prominent dysregulated inflammatory markers. We focus here mainly on IL-1α, IL-1 β, IL-17A and TNF-α and rehearse literature that has previously linked upregulation of these molecules to platelet activation. They are all also known to act as ligands to platelet receptors, which cause outside-in and inside-out platelet signaling See Figure 7 for a simplified diagram of such pathways receptor binding, as well as signaling.

**Figure 7:**
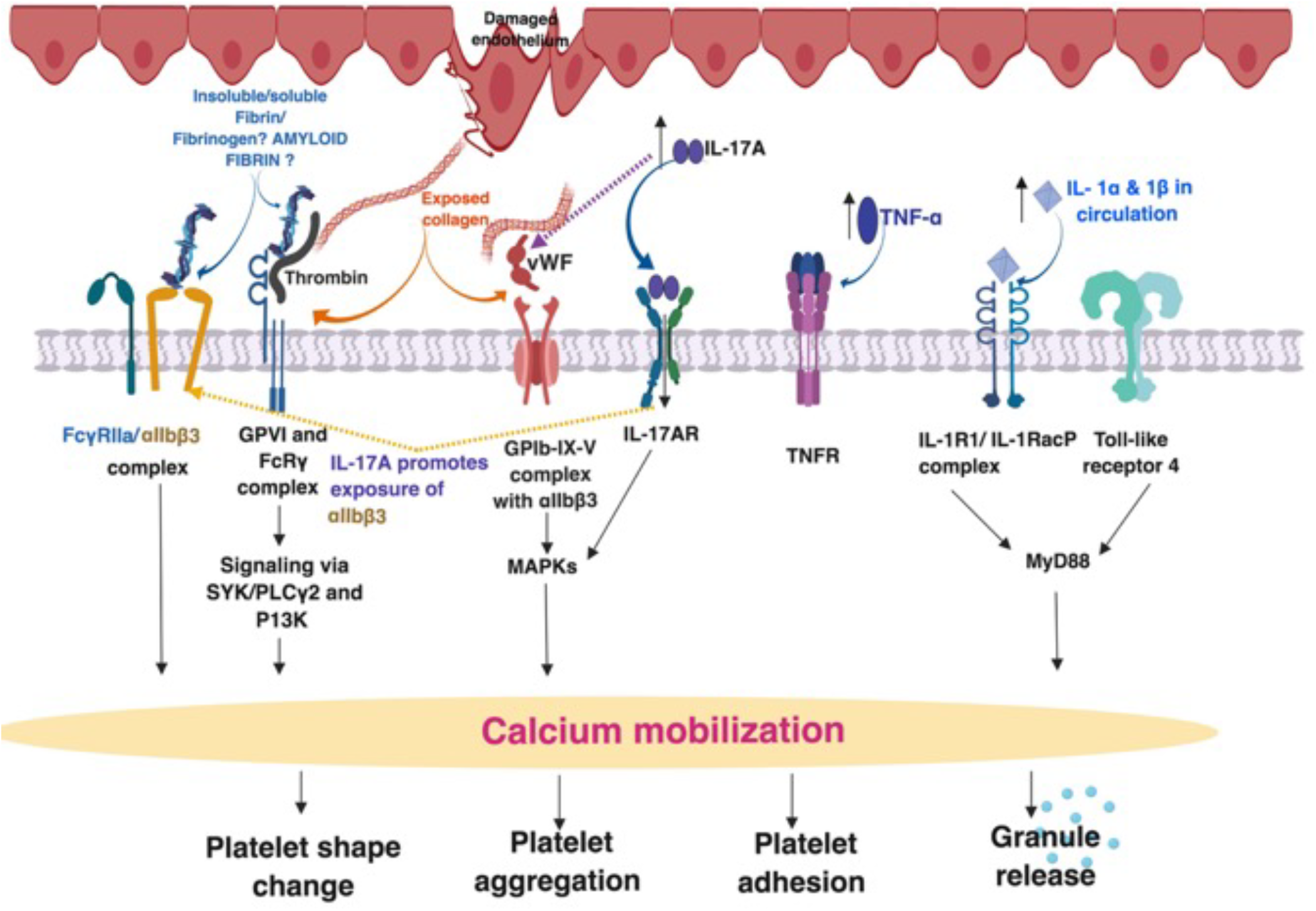
Simplified platelet signaling and receptor activation with main dysregulated molecules IL-1α, IL-1 β, TNF-α, and IL17A. Diagram created using BioRender (https://biorender.com/). When inflammatory molecules are upregulated in circulation, they either cause direct endothelial damage (by binding to receptors on endothelial cells), or they may act as ligands that bind directly to platelet membrane receptors (Olumuyiwa-Akeredolu et al., 2019). When these inflammatory molecules disrupt endothelial cell structure, the endothelial cells release collagen and von Willebrand (vWF). vWF is also a mediator of vascular inflammation (Gragnano et al., 2017), and it binds to exposed collagen and anchors platelets to the subendothelium (Du, 2007), causing platelet aggregation (Xu et al., 2016), and formation of a platelet plug (Jagadapillai et al., 2016). Both collagen and vWF act as platelet receptor ligands, causing platelet outside-in signaling, followed by inside-out signaling. Furthermore, collagen and vWB binding also result in signaling processes that cause a release of stored molecules that are present inside α- and dense granules of platelets, and may also include stored interleukins (e.g. IL-6 and IL-1β); further increasing the concentration of these inflammatory molecules in circulation (Olumuyiwa-Akeredolu et al., 2019). vWF binding is mediated by GpIbα (which is part of the GPIb-IX-V) and integrin αIIbβ3 complex (Bryckaert et al., 2015). This αIIbβ3 receptor also binds fibrinogen and thrombin, and both these molecules and vWF work together to play critical roles in platelet activation and aggregation (Estevez and Du, 2017).

IL-1α, IL-1β, IL17A and TNF-α are all significantly upregulated in our PD sample, and circulating TNF-α, IL-1and IL-17 are also known to stimulate vWF release from damaged endothelial cells (Domingueti et al., 2016; Meiring et al., 2016; Owczarczyk-Saczonek and Placek, 2017). The IL-1 family of ligands and receptors are associated with both acute and chronic inflammation (Gabay et al., 2010; Dinarello, 2011), and IL-1α is an intracellular cytokine involved in various immune responses and inflammatory processes (Schett et al., 2016), and is also known to be upregulated in cardiovascular diseases (Pfeiler et al., 2017). IL-1α has properties of both a cytokine and a transcription factor (Dinarello, 2006), and both IL-1α and IL-1β bind to the IL-1 receptor type 1 (IL-1RI), followed by recruitment of the co-receptor chain, the accessory protein, IL-1RAcP. A complex is formed consisting of IL-1RI, the ligand, IL-1α and the co-receptor (IL-1RAcP). This results in downstream signaling, involving the recruitment of the adaptor protein MyD88 to the Toll-IL-1 receptor domain. Platelets express IL-1R1, as well as Toll-like receptors, and these two receptors are known to be involved in platelet activation, platelet-leucocyte reciprocal activation, and immunopathology (Anselmo et al., 2016). Platelets also signal through the TLR4/MyD88- and cGMP/PKG-dependent pathway (Zhang et al., 2009), causing granule secretion followed by platelet activation and aggregation (Vallance et al., 2017). TNF-α, which is also significantly upregulated in our PD sample, binds to two TNFα receptors that are found on platelets, TNFR1 and TNFR2, resulting in inside-out signaling and platelet (hyper)activation (Pignatelli et al., 2008). Platelets express a receptor for IL-17A, the IL-17R receptor and the cytokine might facilitate their adhesion to damaged endothelium, as well as to other circulating leukocytes, ultimately leading to thrombus formation (Maione et al., 2011). Furthermore, IL-17A facilitates platelet function through the extracellular signal–regulated kinase 2 (ERK2) signaling pathway (part of the MAPK pathway) and causes platelet aggregation (Zhang et al., 2012). IL-17A also promotes the exposure of α_IIb_β_3_ integrin, which provides more ligand binding site for fibrinogen via conformational change, and crosslinks the neighboring activated platelets which results in platelet aggregation (Zhang et al., 2012). These upregulated cytokines in our PD sample therefore could in part be the cause of their hyperactivated platelet ultrastructure shown in Figure 4.

### Amyloid nature of Parkinson’s disease fibrin(ogen)

Previously, we have shown with Thioflavin T that the fibrin(ogen) protein structure in PD can become amyloid in nature, due to mis-folding of the protein (Pretorius et al., 2018c). It is also known that fibrinogen levels in PD is higher than in controls (Wong et al., 2010; Ton et al., 2012). In the current paper, we now include two additional amyloid markers. Our results show enhanced amyloid-fluorescence as assessed by both AmyTracker 480 and 680 and this is confirmed by enhanced Thioflavin T fluorescent in our current PD samples. Our results suggest that in PD clots, fibrinogen polymerises into a form with a greatly increased amount of ß-sheets, reflecting amyloid formation. This reinforces previous data that observed fibrin amyloid in PD using Airyscan (confocal analysis) and Thioflavin T (Pretorius et al., 2018c). This important finding may describe a possible mechanism underlying some of the anomalous clotting formation and coagulopathies occurring in PD. It further emphasizes the systemic nature of PD, demonstrating pathological changes beyond the brain and extending to the circulation. Amyloid fibrin has also been observed in other diseases associated with inflammation and with known hematological abnormalities, including Type 2 Diabetes (Pretorius et al., 2017b; Pretorius et al., 2017c) and Alzheimer’s Disease (Pretorius et al., 2018a). Further, an amyloid state may be induced experimentally by the addition of bacterial membrane products and iron (Pretorius et al., 2018b), as well as products of the acute phase response such as serum amyloid A (Page et al., 2019). These findings imply that the presence of (bacterial) inflammagen molecules, and the inflammatory state more broadly, are conditions that divert fibrinogen polymerization to an amyloid form, and indeed may be overarching (general) features of many chronic, inflammatory diseases (Kell and Pretorius, 2018a).

These results are of particular importance when it is noted that bacterial involvement might play a role in both the development and progression of PD, and specifically, circulating bacterial inflammagens such as LPS have been implicated (Tufekci et al., 2011; De Chiara et al., 2012; Potgieter et al., 2015; Friedland and Chapman, 2017). We have also suggested that LPS may both maintain systemic inflammation, as well as the disease aetiology itself in PD (but also in other inflammatory diseases like type 2 diabetes, pre-eclampsia, sepsis, rheumatoid arthritis and Alzheimer’s disease, where LPS presence has been implicated in the aetiology of the condition) (Kell and Kenny, 2016; Pretorius et al., 2016a; Pretorius et al., 2016b; Pretorius et al., 2017a; Pretorius et al., 2017b; Pretorius et al., 2017c; Kell and Pretorius, 2018b). Indeed in 2018, we showed that LPS from *E. coli* could be identified with fluorescent LPS *E. coli* antibodies in clots of PD, type 2 diabetes and AD (de Waal et al., 2018). There is therefore mounting evidence that PD might have a bacterial involvement, that in part drives the aetiology of the condition. It is recognised that endotoxins (and exotoxins) are among the most potent bacterial inducers of cytokines (Cavaillon, 2018).

### The presence of bacterial inflammagens in Parkinson’s disease

In the current paper, we further investigate <u>*the causative agents*</u> of the amyloid nature of PD fibrin(ogen) and we turned our attention to another prominent bacterium and its inflammagens. *P gingivalis has* long been implicated in PD and periodontitis, and recently its protease (gingipain) was interrogated as a causative agent in AD, where the gingipain proteases was found in brain tissue from patients with AD (Dominy et al., 2019). These researchers also correlated these gingipain quantities within the brain tissue to the extent of tau and amyloid-ẞ pathology. Furthermore, *P. gingivalis* has been found within atherosclerotic tissue of cardiovascular disease patients (Velsko et al., 2014; Olsen and Progulske-Fox, 2015; Atarbashi-Moghadam et al., 2018). Periodontal diseases are a well-known accompaniment to PD (Schwarz et al., 2006; Zlotnik et al., 2015; Kaur et al., 2016; Chen et al., 2017; Chen et al., 2018); however, the direct identification of *P. gingivalis* or its molecular signatures in circulation and/or brain tissue of PD patients has not previously been made.

Previous studies conducted on fibrinogen and plasma have shown that Rgp and Kgp increase thrombin time when compared to control samples (Imamura et al., 1995). Furthermore, the activation of other coagulation factors by gingipains have been established, including factor IX, X and prothrombin prothrombin (Imamura et al., 1997; Imamura et al., 2001). Based on these observations, there seems to be a major disruption in the homeostatic control of the coagulation system/cascade when gingipain proteases are present. Here, we show that RgpA protease produced by *P. gingivalis* is present in PPP clots from our PD sample blood using polyclonal antibodies. We also confirmed that in control PPP clots, little to no signal was noted, but that we could detect RgpA with its fluorescent antibody in control clots after addition of the recombinant protease to healthy PPP. In addition, we used a fluorescent purified fibrinogen model to show that LPS from *P. gingivalis* can cause hypercoagulability and that RgpA could hydrolyse fibrin(ogen) to such an extent that healthy clot formation is impaired. However, when both *P. gingivalis* LPS and RgpA are co-incubated, abnormal (hyperclottable) fibrin(ogen) is still visible. These results are in line with our finding that in PD clots are more dense and hyperclottable (Pretorius et al., 2018c). It also supports our current TEG results that showed a hyperclottable clot phenotype.

We conclude by suggesting that our results strongly support a systemic inflammatory and hypercoagulable aetiology fueled by a bacterial presence, and serves as a preliminary study showing a role of *P. gingivalis* LPS and gingipain protease in abnormal blood clotting observed in our PD sample. The next step would be to identify the extent to which this bacterium might contribute to Parkinson’s pathology or if there are any specific links, e.g. a link with the presence of α-synuclein and auto/xenophagy (El-Awady et al., 2015; Cerri and Blandini, 2018). Furthermore, our finding that gingipain antibody signal was detected in clots from our PD samples but not the control emphasizes the possibility of this bacterium having a role in PD pathology. We have discussed research that pointed to the fact that bacteria, more generally, are implicated in PD aetiology, and here we note the possible involvement of *P. gingivalis*, specifically. Taking these findings in both a neurological and cardiovascular context, it is plausible to believe that the entry, dissemination and infection of this bacterium and its virulent machinery in a systemic manner may be an aetiological and/or driving factor for disease worth investigating.

## LIST OF ABBREVIATIONS

IFN-α: Interferon-alpha
IL-10: Interleukin-10
IL-13: Interleukin-13
IL-4: Interleukin-4
E-Selectin: E-Selectin
GM-CSF: Granulocyte-macrophage colony-stimulating factor
IL-1γ: Interferon-gamma
IL-1α: Interleukin-1 alpha
IL-1β: Interleukin-1 beta
IL-12p70: Interleukin-12p70
IL-17A: Interleukin-17A
IL-6: Interleukin-6
IL-8: Interleukin-8
IP-10: Interferon gamma-induced protein-10
MCP-1: Monocyte chemoattractant protein-1
MIP-1α: macrophage inflammatory protein-1 alpha
MIP-1β: macrophage inflammatory protein-1 beta
P-Selectin: P-Selectin
sICAM-1: Soluble intercellular adhesion molecule-1
TNF-α: Tumor necrosis factor-alpha
RgpA: Recombinant gingipain R1 protease
ERK2: Extracellular signal–regulated kinase 2

## DISCLOSURES AND ACKNOWLEDGEMENTS

### Author contribution statement

BA: patient blood collection and preparation of blood samples; TEG and 20-plex analysis, statistics; TAN: statistics and editing of paper; JMN: gingipain experiments; MJP: amyloid assay and editing of the paper; TR: all correlation analysis and plots; EP: study leader, writing of the paper and co-corresponding author; JC: clinician; DBK: edited the paper and co-corresponding author. All authors reviewed the manuscript.

## Acknowledgements

We thank the Biotechnology and Biological Sciences Research Council (grant BB/L025752/1) as well as the National Research Foundation (NRF) of South Africa (91548: Competitive Program) and the Medical Research Council of South Africa (MRC) (Self-Initiated Research Program) for supporting this collaboration. The funders had no role in study design, data collection and analysis, decision to publish, or preparation of the manuscript.

